# Host porphobilinogen deaminase deficiency confers malaria resistance in *Plasmodium chabaudi* but not in *Plasmodium berghei* or *Plasmodium falciparum* during intraerythrocytic growth

**DOI:** 10.1101/589242

**Authors:** Cilly Bernardette Schnider, Hao Yang, Lora Starrs, Anna Ehmann, Farid Rahimi, Elena Di Pierro, Giovanna Graziadei, Kathryn Matthews, Tania De Koning-Ward, Denis C. Bauer, Simon J. Foote, Gaetan Burgio, Brendan J. McMorran

## Abstract

An important component in host resistance to malaria infection are inherited mutations that give rise to abnormalities and deficiencies in erythrocyte proteins and enzymes. Understanding how such mutations confer protection against the disease may be useful for developing new treatment strategies. A mouse ENU-induced mutagenesis screen for novel malaria resistance-conferring mutations identified a novel nonsense mutation in the gene encoding porphobilinogen deaminase (PBGD) in mice, denoted here as *Pbgd*^*MRI58155*^. Heterozygote *Pbgd*^*MRI58155*^ mice exhibited approximately 50% reduction in cellular PBGD activity in both mature erythrocytes and reticulocytes, although enzyme activity was approximately 10 times higher in reticulocytes than erythrocytes. When challenged with blood-stage *P. chabaudi*, which preferentially infects erythrocytes, heterozygote mice showed a modest but significant resistance to infection, including reduced parasite growth. A series of assays conducted to investigate the mechanism of resistance indicated that mutant erythrocyte invasion by *P. chabaudi* was normal, but that following intraerythrocytic establishment a significantly greater proportions of parasites died and therefore affected their ability to propagate. The *Plasmodium* resistance phenotype was not recapitulated in *Pbgd*-deficient mice infected with *P. berghei*, which prefers reticulocytes, or when *P. falciparum* was cultured in erythrocytes from patients with acute intermittent porphyria (AIP), which had modest (20-50%) reduced levels of PBGD. Furthermore, the growth of *Pbgd*-null *P. falciparum* and *Pbgd*-null *P. berghei* parasites, which grew at the same rate as their wild-type counterparts in normal cells, were not affected by the PBGD-deficient background of the AIP erythrocytes or *Pbgd*-deficient mice. Our results confirm the dispensability of parasite PBGD for *P. berghei* infection and intraerythrocytic growth of *P. falciparum*, but for the first time identify a requirement for host erythrocyte PBGD by *P. chabaudi* during *in vivo* blood stage infection.

**IMPORTANCE:** The causative agent of malaria, *Plasmodium*, adopts a parasitic lifestyle during erythrocyte infection, and as such relies on host cell factors for its survival and growth. Host-encoded mutations that alter the availability of these factors confer disease resistance, including several well-known genetic erythrocyte abnormalities that have arisen due to the historical evolutionary pressure of malaria. This study identified in mice a novel malaria resistance-conferring host mutation in the heme biosynthesis enzyme, porphobilinogen deaminase (PBGD), and compared the relative requirements by *Plasmodium* for the host versus parasite-encoded forms of PBGD in both *in vivo* and *in vitro* settings. The findings demonstrated that parasite PBGD was dispensable, but that the host enzyme was important specifically during *in vivo* infection by *P. chabaudi*, and collectively suggest that *Plasmodium* requires a certain threshold of the enzyme to sustain its intraerythrocytic growth. *Plasmodium* may therefore be vulnerable to other interventions that limit host PBGD activity.

## INTRODUCTION

Symptomatic and lethal stages of malaria occur when *Plasmodium* parasites invade and replicate within erythrocytes. The ongoing relationship between parasite and host, spanning primordial vertebrates and human evolutionary history, has exerted significant selective pressures on human populations. This has manifested as polymorphisms and mutations that produce altered erythrocytic proteins, many of which are associated with host resistance to infection. These include hemoglobinopathies, and deficiencies in cytoskeletal proteins and various enzymes that are highly expressed in erythrocytes; they act either directly by impeding parasite invasion or growth, or indirectly on host immune responses and parasite clearance [1; 2; 3; 4]. By virtue of its parasitic lifestyle during the blood-stage of the disease, *Plasmodium* also scavenges erythrocyte enzymes, including protein kinases, cellular redox regulators, and those involved in heme biosynthesis [5; 6; 7; 8; 9]. Depriving parasites of such host enzymes may contribute to resistance to *Plasmodium* infection and facilitate identifying novel antimalarial therapeutics as reported in model systems [10; 11].

Heme is an essential cofactor in many proteins and enzymes, for example in hemoglobin as the carrier of oxygen. The canonical heme biosynthetic pathway uses eight enzymes, which catalyze conversion of glycine and succinyl-CoA into protoporphyrin, and subsequent Fe^2+^ incorporation. Heme biosynthesis is essential in humans and complete deficiency of any of the synthetic enzymes is lethal. However, mutations that reduce enzyme production or activity are known and cause porphyria. Characteristic symptoms of porphyria may include mild to severe skin photosensitivity, liver malfunction and neurological problems, and are caused by build-up of toxic porphyrin precursors and their derivatives. Specific symptoms and disease severity depend on the composition and levels of these molecules, which in turn are determined by enzyme activity and responsible mutations.

*Plasmodium* has a canonical heme biosynthesis pathway [12]. Interestingly, essentiality of the pathway differs depending on the parasite life-cycle stage. *P. berghei* made genetically deficient in either the first (δ-aminolevulinic acid synthase; ALAS) or last (ferrochelatase; FC) enzyme of the pathway are not viable in the mosquito stages, but propagate in the mouse bloodstream and cause symptoms that are indistinguishable from those by wild-type parasites [12; 13]. Another study in *P. berghei* has reported a similar requirement for porphobilinogen deaminase (PBGD) in mosquito but not erythrocyte stages of growth [14]. Similarly, *P. falciparum* carrying deletions in FC, PBGD or another pathway enzyme, coproporphyrinogen oxidase grow normally when cultured in human erythrocytes [12; 15].

How these enzyme-deficient parasite lines obtain heme is topic of some controversy. Based on the observation that no heme was synthesized in an FC-deficient *P. falciparum* [12], acquisition directly from the degradation of host hemoglobin was suggested a likely source [12]. However, co-opting of the host heme-biosynthetic enzymes to participate in *de novo* synthesis has been demonstrated in other studies [5; 9; 16]. Notably, although heme biosynthesis is restricted to erythroid progenitors, mature erythrocytes contain detectable amounts of active enzymes [17; 18; 19] including PBGD, which is biosynthetically active in reticulocytes and mature erythrocytes [15; 20].

Previously, we showed that the growth of both *P. chabaudi* and *P. falciparum* was significantly perturbed in mouse and human erythrocytes with reduced levels of FC (erythropoietic protoporphyric erythrocytes). In contrast, *P. falciparum* grew normally in X-linked dominant protoporphyric erythrocytes, which have normal levels of FC, but elevated porphyrin precursors [21]. In addition, treatment with FC substrate analogues that specifically inhibit the enzyme killed cultured blood stage *P. falciparum*, and inclusion of excess enzyme substrate rescued the effect [11; 21]. Collectively, this indicated that *Plasmodium* has a specific requirement for host FC to sustain its intraerythrocytic growth in erythrocytes. Here we identified a novel mutation in the gene encoding PBGD in mice that reduced erythrocytic enzyme activity and compared the growth requirements of *Plasmodium* for host versus parasite-encoded PBGD in both *in vitro* and *in vivo* settings of blood stage infection.

## RESULTS

In a large-scale ENU mutagenesis screen for abnormal red blood cell count, we discovered a mouse, denoted as MRI58155, which displayed microcytosis accompanied by moderately elevated reticulocyte counts, reduced hematocrit and total hemoglobin, but with normal mean erythrocytic hemoglobin levels. We confirmed that the phenotype was heritable after crossing the MRI58155 G1 animal with SJL/J mice and observing that the G1 microcytosis phenotype was passed on to approximately half the offspring. Intercrossing two affected G2 mice produced the wild-type phenotype in approximately one third of the progeny and the G1 founder phenotype in the other two thirds (Table S1). Collectively this indicated a fully penetrant dominant inheritance of the microcytotic phenotype (i.e., heterozygous for the MRI58155 allele), but embryonic lethality in homozygotes.

Whole-exome sequencing of two heterozygous G3 mice and subsequent segregation analysis of mutations in candidate genes identified a single base substitution (A to G) in chromosome 9 (Chr9 nucleotide position 44341074, mm10), which is located in the murine *Pbgd* gene (NM_013551). This mutation designated *Pbgd*^*MRI58155*^ is predicted to disrupt splicing of exon 6 and result in a truncated protein comprising 90 amino acids; this theoretical protein lacks all conserved residues and domains required for catalytic function of PBGD (Figure 1A). The mutation completely segregated with the microcytic phenotype over more than 10 generations of breeding, excluding the possibility that other mutations were involved in the phenotype. No detectable differences in PBGD protein levels were seen in extracts from spleen, liver, bone marrow, or erythrocytes from heterozygous versus wild-type mice and the predicted 20 KD truncated was not observed (data not shown). However, PBGD enzyme activity, measured using a specific and sensitive assay, was approximately 50% lower in heterozygous versus wild-type purified mature erythrocytes (3.38 ± 1.80 versus 6.53 ± 2.14 nkat/L; *p* = 0.048) (Figure 1B). We also separately purified and assayed PBGD activity in reticulocytes. Activity in the mutant reticulocytes was also relatively lower than in wild-type reticulocytes (22.1 ± 14.3 versus 60.0 ± 36.3 nkat/L; *p* = 0.08) (Figure 1B), although, notably, the reticulocyte values were approximately 10 times higher than respective erythrocyte values. Urine from heterozygous mice appeared slightly pink, indicative of excretion of excess porphoryrin synthetic precursors. The heterozygous mice also exhibited a modest splenomegaly, but normal liver size and normal iron content in both organs; serum cytokine levels were also similar to those in wild-type mice (Tables S2 and S3). Erythrocyte life span (Figure S1A) and osmotic fragility (Figure S1B) were also unaffected by the mutation. In addition, production of erythroid progenitors was unaffected (Figure S1C). No homozygous animals were generated from intercrossing more than different 10 pairs of *Pbgd*^*MRI58155*^ heterozygotes, indicating the homozygous *Pbgd*^*MRI58155*^ mutation was probably embryonic lethal.

**Figure 1.**
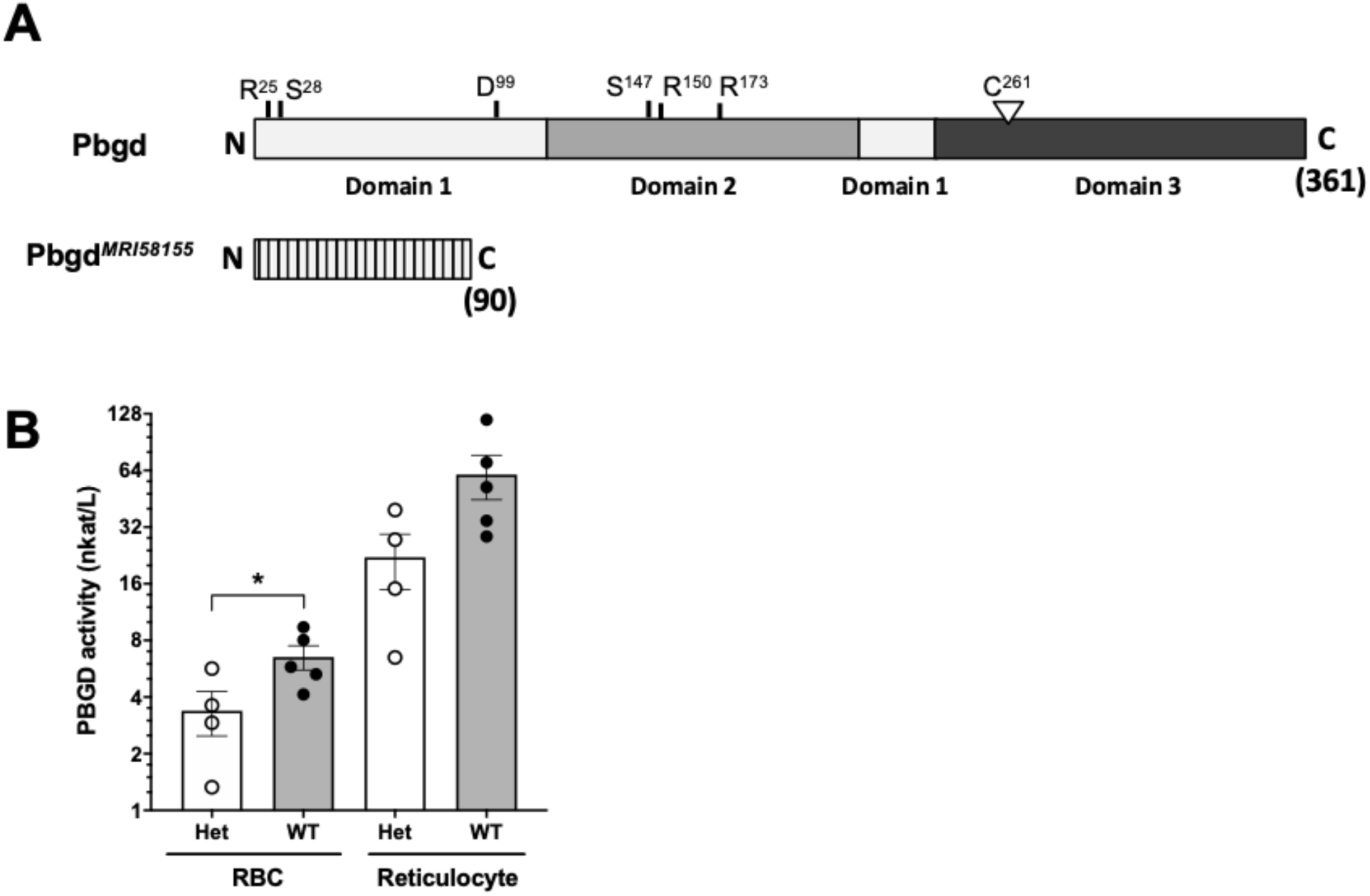
Analysis of the *Pbgd*^*MRI58155*^ mutation. **(A)** Schematic of the full-length 361 residue murine PBGD protein, including locations of the three domains and active site residues, R^25^ and S^28^ (sulfate-binding), D^99^ (catalytic), S^147^, R^150^ and R^173^ (pyrrole-cofactor-interacting) and C^261^ (pyrrole-cofactor-binding), based on a solved human PBGD crystal structure; Gill et al 2009). The predicted 90 residue PBGD^MRI58155^ protein is shown below. Its lacks the sequences containing the catalytic domain and cofactor binding sites. (**B**) Measurement of PBGD enzyme activity in whole blood samples from WT and Het mice using HPLC. Data represent the mean (box) and values from four mice per group, each assayed two times. * *p* < 0.05, calculated using a two-tailed *t-*test assuming equal variance.

To investigate if the *Pbgd*^*MRI58155*^ mutation affects the host resistance to malarial infection, cohorts of heterozygous and wild-type mice were inoculated with blood stage *P. chabaudi adami* DS, which is typically lethal to this parental mouse strain (SJL/J). We observed an altered course of infection in heterozygotes compared to wild-types. Parasitemias were significantly lower in the heterozygotes between days seven and ten in female mice (Figure 2A), and days nine and ten in male mice (Figure 2B). A modest but highly significant survival advantage was also observed for female heterozygotes (*p* = 0.0016); 90% of these animals survived for at least one day longer than their wild-type littermates, although 12 of 13 heterozygous mice eventually succumbed (Figure 2C). Two of 21 heterozygous male mice survived the infection, but this was not statistically significant (Figure 2D). We investigated whether the reduced parasitemia observed in the heterozygous mice was due to compromised merozoite invasion of erythrocytes. To assess this we used an *in vivo* erythrocyte-tracking assay (IVET) [22], where a mixture of heterozygous and wild-type erythrocytes, each labelled with a different fluorescent dye, was transfused into recipient wild-type mice infected with *P. chabaudi* (at the point of parasite schizogony), and the relative proportions of donor cells that became infected were determined over time. There were no differences between the proportions infected heterozygous and wild-type cells measured between 30 min and 20 h after cell transfer. After 39 h the parasitemia increased relative to the earlier timepoints, reflecting a second schizogony and infection cycle, but there were still no apparent differences between the donor cells (Figure 2E). These results indicate that the *P. chabaudi* merozoites invade *Pbgd*^*MRI58155*^ heterozygous and wild-type erythrocytes at similar rates. We also investigated if *P. chabaudi* intraerythrocytic growth was impaired by PBGD insufficiency. Dead or dying intraerythrocytic *P. chabaudi* were identified using an adapted terminal deoxynucleotidyl transferase dUTP nick end labeling (TUNEL) assay [23]. The mean percentages (± SEM) of TUNEL-labeled parasites measured in mouse blood samples collected eight and nine days after *P. chabaudi* infection were 9.0 ± 1.0 and 12.6 ± 1.2 in wild-type mice and 10.4 ± 1.4 and 15.8 ± 1.7 in heterozygous mice; the heterozygous values were significantly greater than those of the wild-type mice at both time points (*p* <0.05; Figure 2F). This finding suggests that in heterozygous erythrocytes, a greater proportion of parasites were dying, which in turn may reduce production of new, viable parasites and thereby provides an explanation for the observed reduction in parasitemia and improved survival.

**Figure 2:**
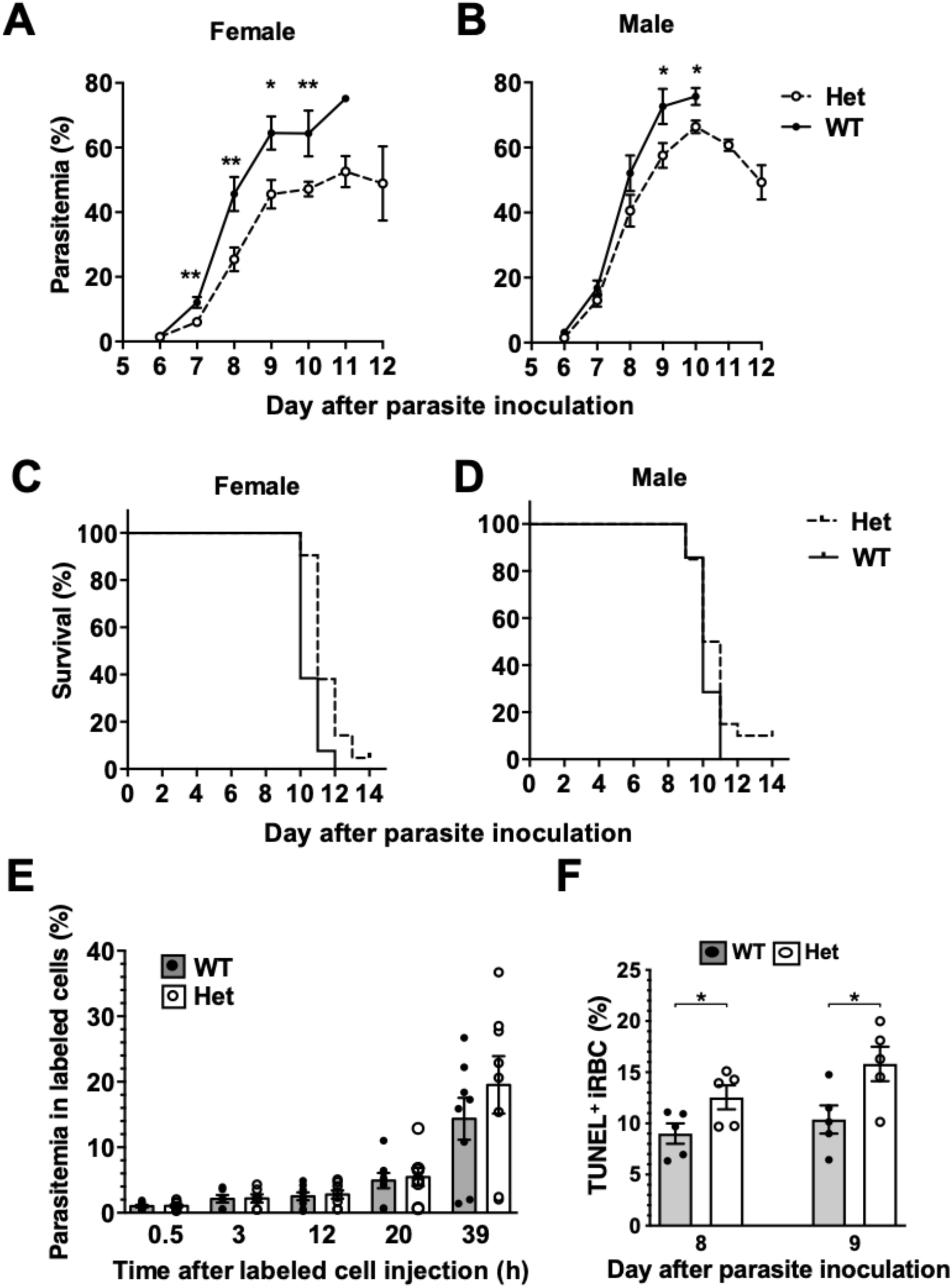
Mice with the *Pbgd*^*MRI58155*^ mutation display an increased resistance to infection with *P. chabaudi*. **(A and B)** Parasite growth and **(C and D)** survival in *Pbgd*^*MRI58155*^ heterozygotes (Het) and wild-type littermates (WT) infected with *P. chabaudi* (1×10^4^ parasites; 13-21 mice per group). **(E)** Parasitemia measured at various timepoints in labeled donor RBC following transfusion into *P. chabaudi* infected recipient mice (eight days after parasite inoculation). Donor RBC were from *Pbgd*^*MRI58155*^ heterozygotes (Het) or wild-type littermates (WT) (8 recipient mice per group). **(F)** Proportions of TUNEL-labeled parasite-infected erythrocytes in *Pbgd*^*MRI58155*^ heterozygotes (Het) or wild-type littermates (WT) infected with *P. chabaudi* (five per group). Error bars represent SEM. * *p* < 0.05, ** *p* < 0.01, calculated using a two-tailed *t*-test assuming equal variance. Log-rank (Mantel-Cox test) for differences in survival yielded *p* = 0.0016 (female group) and *p* = 0.16 (male).

The *Pbgd*^*MRI58155*^ heterozygous mice were also challenged with another murine malarial parasite, *P. berghei* ANKA (Pb ANKA), which preferentially infects reticulocytes and in the SJL/J mouse strain causes severe anemia without any cerebral malaria symptoms. We also generated a *P. berghei* parasite line in which the parasite *Pbgd* (PbA_060800) was deleted (PbA_Pbgd, Figure S2). There were no differences in the parasite growth rates between the heterozygous and wild-type mice infected with wild-type parasites, and the *Pbgd* knockout and wild-type parasites propagated equivalently in both mouse strains (Figure 3). There were also no differences in the disease course or susceptibility over the period of the infection (17 days); mice were ethically euthanized after this time. Collectively these data indicate that the *Pbgd*^*MRI58155*^ mutation conferred resistance to *P. chabaudi* infection but not *P. berghei* infection, and that the parasite-encoded PBGD was dispensable for *P. berghei* infection irrespective of the host PBGD sufficiency status.

**Figure 3.**
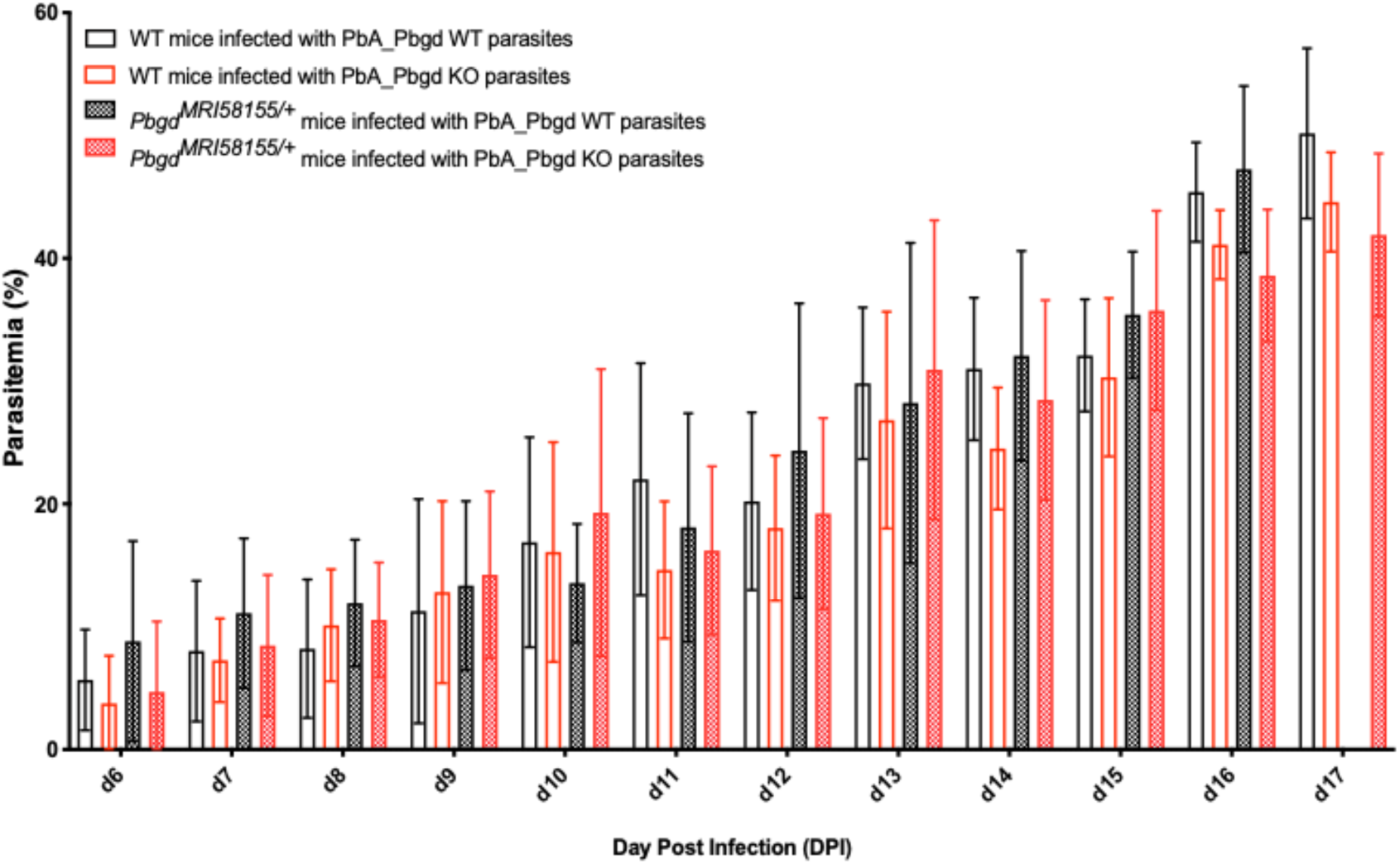
Parasite growth analysis of wild-type and *Pbgd*-knockout *P. berghei* in wild-type and *Pbgd*^*MRI58155/+*^ deficient mice. Blood parasitemia determined in WT and *Pbgd*^*MRI58155*^ heterozygous (Het) mice after infection with either *Pbgd*-knockout *P. berghei* (*PbA_Pbgd* KO) or parental wild-type (WT). For parasite lines the starting inoculum was 1×104 parasites. The data represent the mean (+/- SD) for 8 to 15 mice per group. Data per group). growth in infected WT and *Pbgd*^*MRI58155*^ heterozygous mice. There were no statistically significant differences comparing the four different parasite and animal combinations on each day of the infection using two-way ANOVA.

PBGD deficiency in humans can manifest as AIP (MIM 176000). AIP is characterized by occasional, acute episodes of gastrointestinal and neuropathic symptoms; between episodes, the patient is healthy. Acute attacks result in high urinary excretion of the heme precursors, especially δ-aminolevulinic acid and porphobilinogen [24]. We obtained blood samples from four AIP patients (A – D), all with the same confirmed *PBGD* gene mutation (PBGD c.[913 G>C];[=]). This mutation has not been reported previously. It is predicted to cause a splice site mutation and premature termination of translation, resulting in a carboxyl-terminal deletion (His305-Ter362) and an inactive protein. PBGD enzyme activity levels measured in whole blood samples from these patients were between 50 and 80% of normal levels (Table S4). All of the patients reported a typical history of episodes of acute symptoms with elevated urinary δ-aminolevulinic acid and porphobilinogen levels, and two were symptomatic at the time of blood collection.

Isolated erythrocytes from these AIP patients as well as two healthy individuals, one collected in Italy (Normal Italy) and one from an Australian Red Cross Blood Service red cell donor (Normal ARC), were infected with growth-stage-synchronized *P. falciparum* (pigmented trophozoites and schizonts). Cultures were incubated for up to 72 h and sampled at different time points to count the number of parasitized cells and compare parasite propagation over time. Following the first 24 h of culturing, after parasites had egressed from donor erythrocytes and invaded the surrounding test cells, we observed no differences in parasitemia fold-change between the AIP and healthy control cells. After 48 h, parasites had developed from early ring stage into mature trophozoite and schizont forms of the parasite, and similar proportions were observed between the AIP and control cells. After 72 h, parasites had replicated in the test erythrocytes and undergone a further cycle of schizogony and cell invasion; the parasitemia fold changes in all of the control and AIP samples were also similar at this time point (Figure 4). In the same experiments we also infected these control and AIP cells with a *P. falciparum* clone carrying a genetic disruption of the parasite *Pbgd* homolog (PF3D7_1209600); this was shown previously to grow as well as its wild-type parent strain (3D7) in normal human erythrocytes [15]. The *Pbgd* deleted line grew similarly to the wild-type line in all of the erythrocyte samples, and no differences in parasitemia fold change were observed after 24 or 72 h of culture (Figure 4). Collectively, these results indicated that neither the invasion or intraerythrocytic growth of both wild-type and *Pbgd* deleted *P. falciparum* were sensitive to the AIP host cell PBGD insufficiency.

**Figure 4.**
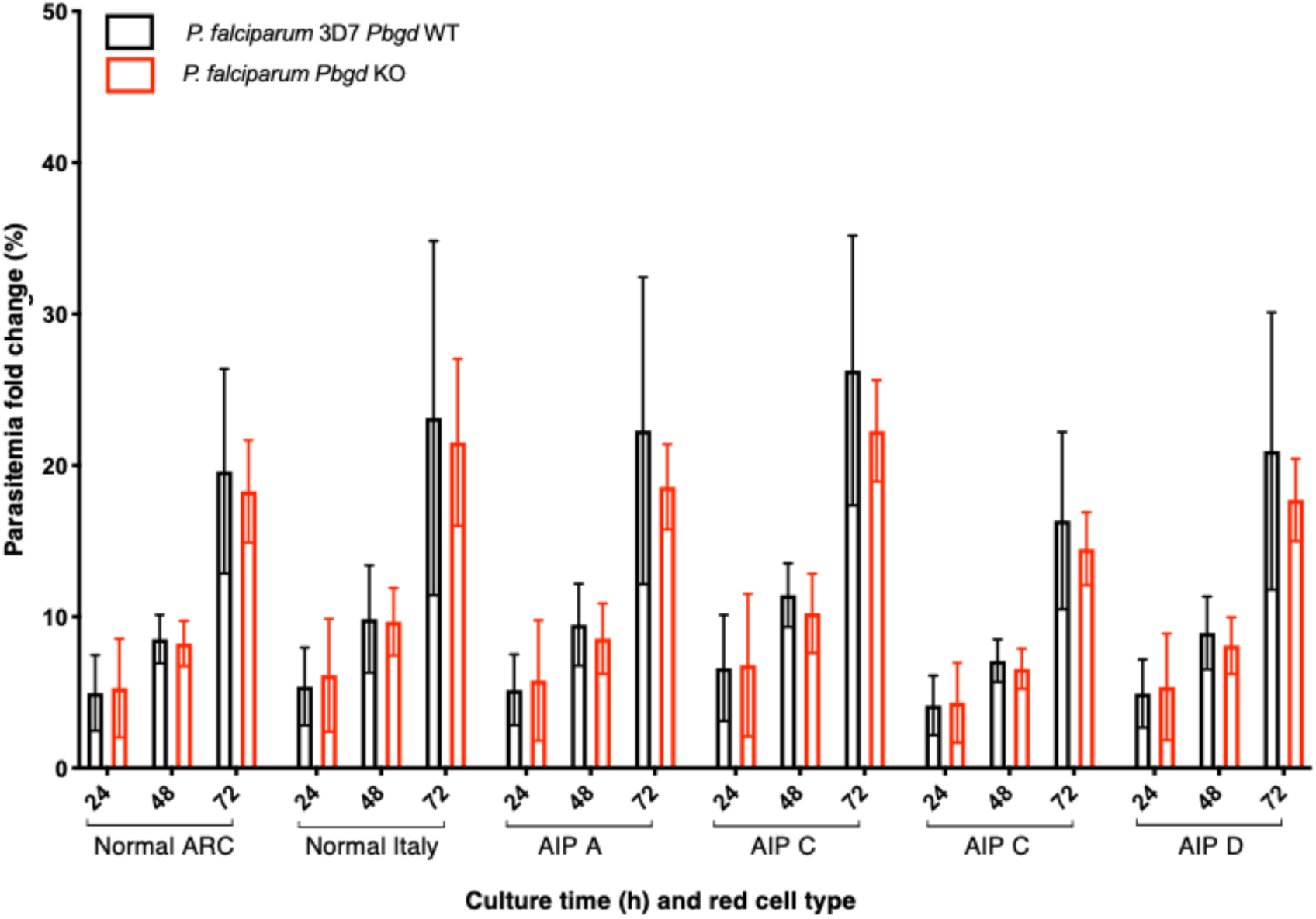
Parasite growth analysis of wild-type and *Pbgd*-knockout *P. falciparum* in erythrocytes from normal and AIP individuals. Parasite growth following inoculation of wild-type (*Pbgd* WT) or *Pbgd*-knockout (*Pbgd* KO) *P. falciparum* into erythrocytes isolated from healthy individuals (Normal ARC and Normal Italy) and from four AIP individuals (A-D; see Table S4 for biochemical and PBGD activity characteristics). The parasites were cultured for up to 72 h. Data represent the mean fold changes (+- SD) for two independent growth assay experiments, except for AIP B which was assayed once. There were no statistically significant differences comparing between the Normal and AIP samples, or between the two parasite strains using two-way ANOVA.

## DISCUSSION

Here we investigated if malarial parasites require the heme biosynthetic enzyme PBGD for intraerythrocytic growth and infection. This idea was motivated by previous observations that other host heme biosynthetic enzymes as well as precursors of the pathway are co-opted by the parasite [5; 9; 11; 16; 21]. To do this, we used a combination of genetic models of PBGD insufficiency and deficiency in mouse, human and *Plasmodium* to compare and contrast parasite growth and the mechanisms of host resistance to malarial infection.

In mice, PBGD insufficiency was generated through the identification and characterization of a novel ENU-induced mutation in the murine *Pbgd* gene (*Pbgd*^*MRI58155*^). Mice heterozygous for this mutation displayed a relatively normal phenotype, distinguished by microcytosis and a modest, but significant decrease in PBGD activity, although they exhibited no overt symptoms characteristic of AIP. Previous studies have shown that a profound deficiency of PBGD in mice is necessary to mimic human AIP [25]. The protein expressed by the *Pbgd*^*MRI58155*^ allele is predicted to be truncated and completely inactive. Consistent with this prediction, we never obtained homozygous pups and demonstrated approximate 50% reductions PBGD enzyme activity in both mature reticulocyte-depleted erythrocytes and in purified reticulocytes. The most prominent consequence of the *Pbgd*^*MRI58155*^ mutation, and indeed the phenotypic basis of its identity, was a modest microcytic anemia, although erythrocyte number and erythrocytic hemoglobin concentrations were normal. Reduced biosynthesis of heme in erythroid progenitors can often lead to generation of microcytic anemia, although this is not characteristic of human AIP, and is not a reported effect in mouse lines with the *Pbgd-*null or hypomorphic mutations [25]. However, microcytic erythrocytes have been observed in cats with *Pbgd* mutations displaying AIP symptoms [26]. Overall, the *Pbgd*^*MRI58155*^ allele results in a phenotype consistent with a mild, asymptomatic form of AIP.

We found that the propagation of the murine malarial parasite *P. chabaudi* was reduced in the *Pbgd*^*MRI58155*^ heterozygous mice and survival to infection was improved. Our data suggests the most likely reason for the resistance is the compromised ability of *P. chabaudi* to survive within the mutant erythrocytes. We specifically excluded other possible explanations for the resistance, including merozoite invasion, immune responses, iron status and erythrocyte structure and function. However, another murine-specific species *P. berghei* grew normally in the *Pbgd*^*MRI58155*^ mice, as did *P. falciparum* cultured in human AIP (PBGD-insufficient) erythrocytes. The growth and infectivity of *P. falciparum* and *P. berghei* were also compared with genetically modified lines in which the respective parasite *Pbgd* loci were deleted (including a new *Pbgd*-null *P. berghei* line generated in this study). Each *Pbgd*-deleted species grew as well as their wild-type counterpart when introduced into the normal and AIP erythrocytes, or the wild-type and *Pbgd*^*MRI58155*^ mice, respectively. Collectively, these results suggest that parasite PBGD is completely dispensable for blood stage growth of *P. falciparum* and *P. berghei*, confirming previous studies [14; 15] and, at least under the conditions used here to grow the parasites, that host erythrocyte PBGD is also not important. In order to understand why *P. chabaudi* but not *P. berghei* exhibits reduced growth in the *Pbgd*^*MRI58155*^ mouse, an important distinction is the approximate 10-fold lower PBGD activity measured in the erythrocytes versus reticulocytes. *P. chabaudi* and *P. berghei* have respective tropisms for these cells, so the former must survive in a lower PBGD environment and may therefore be more sensitive to the added scarcity of PBGD in the mutant cells. In other words, the parasite may require a certain threshold level of host PBGD to efficiently develop and propagate within the erythrocyte. It is also possible that the mutation has other indirect effects on *P. chabaudi* but not *P. berghei* or that these species have differences in their acquisition or requirement for heme. The relatively mild decreased PBGD activities in the AIP erythrocytes may be insufficient to affect *P. falciparum* (also an erythrocyte-preferring species) under culture conditions.

In conclusion, our data suggest that erythrocyte-stage *Plasmodium* is sensitive to profound but not mild deficiencies in erythrocyte PBGD enzyme activity and speculate other strategies that acutely reduce or temporally block PBGD could be effective at preventing parasite growth.

## METHODS

### Ethics statement

Blood sampling and experimental procedures involving patients were performed in accordance with the 1983 revision of the Declaration of Helsinki. The study was approved by the Ethical Committee of Fondazione IRCCS Ca’ Granda, Ospedale Maggiore Policlinico, Milan, Italy (Project Number 246_2015) and The Australian National University Human Research Ethics Committee (2014/765). Written informed consent was obtained for each subject prior to study participation.

All procedures involving animals conformed to the *Australian code of practice for the care and use of animals for scientific purposes* guidelines by the Australian National Health and Medical Research Council and were approved by Macquarie University Animal Ethics Committee (Project Numbers: ARA 2012/017 and 2012/019), the Australian National University Animal Experimentation Ethics Committee (2014/53) and Deakin University animal ethics committee (Project Number G37-2013).

### Animals

Mice were bred and housed under specific pathogen-free conditions. Mice were housed under controlled temperature (21°C) with a 12:12 h light–dark cycle and fed a diet of Rat and Mouse Premium Breeder chow (Gordons Specialty Feeds, Glen Forest, WA, Australia). Seven-week-old SJL/J male mice were injected intraperitoneally (IP) with 150 mg/kg of *N*-ethyl-*N*-nitrosourea (ENU) three times at one-week intervals. Mutagenized G0 mice were crossed with background SJL/J mice. G1s were bled after seven weeks for blood analysis. The G1 58155 founder displayed a mean corpuscular volume that was two standard deviations smaller than average for the colony; it was crossed with SJL/J and resulting G2s were used in further experiments. For each experimental procedure, heterozygous *Pbgd*^*MRI58155*^ mice were compared to their wild-type littermates and housed together, up to five animals per cage (Green Line IVC, Tecniplast, PA). Sample sizes were chosen based on statistical power calculations to limit the number of animals used. For generating *P. berghei Pbgd-*knockout parasites, 6-8 week-old BALB/c mice were used.

### Mouse phenotyping

Complete blood cell counts were obtained using an ADVIA® 2120 hematology system. Erythroid progenitors were examined by staining the bone-marrow-derived cells from mutant and wild-type mice with anti-TER119 and anti-CD44 (eBioscience). Cells were analyzed on a BD FACSARIA™-II flow cytometer, recording 10,000 events per sample. Populations were gated as described [27]. Cytokine Flexset assays (BD Bioscience) were used to measure plasma cytokine concentrations according to the manufacturer’s instructions.

Erythrocyte life span was determined by following intravenous administration of 1 mg of NHS-biotin-ester dissolved in 0.9% (w/v) saline, and blood sampling by tail-snipping each day for seven days, and staining with anti-TER119 and anti-CD71 (eBioscience). Cells were suspended in phosphate-buffered saline (PBS; 10 mM sodium phosphate, 2.7 mM KCl, 140 mM NaCl, pH 7.4) containing 1% (w/v) BSA solution containing 10^8^ counting beads/μL, and the number of TER119^+^ CD71^−^ cells calculated per bead.

Hepatic and splenic iron levels were measured in tissues dried at 45 °C for 48 h and then digested in a 10% hydrochloric acid/10% trichloroacetic acid (TCA) solution for 48 h at 65 °C, and values normalized according to tissue dry weight. Samples were centrifuged (10000×*g* for 5 min); 200 μL of supernatant was added to 1 mL of 1,10-phenanthroline monohydrate solution and incubated for 15 min at ∼23 °C. After incubation, absorbance was measured at λ = 508 nm.

Erythrocytic osmotic fragility was determined by incubating blood samples diluted 1:100 in NaCl solutions ranging from 0 to 160 mM. After 30 min at 37 °C, samples were centrifuged and absorbance of the supernatant measured at λ = 545 nm. Percentage lysis was calculated by comparison to 100% lysis in pure water.

### Whole-exome sequencing

DNA from two MRI58155 mice was isolated using a Qiagen DNeasy Blood and Tissue Kit (Hilden, Germany). DNA (≥10 μg) was prepared for paired-end genomic library sequencing using a kit from Illumina (San Diego, CA). Exomes were enriched using the Agilent SureSelect kit. Samples were sequenced using a Hiseq 2000 platform. Variants were identified and filtered as described [28]. Briefly, we mapped the short sequence tags (reads) on the mouse genome (mm10/NCBI38) using BWA V0.61 [29] and BOWTIE2 [30]. Nucleotide variants were identified using SAMTOOLS V0.1.19 [31] and GATK [32]. Through our variant-filtration process, we retained those variants that are “common” and “private” to the two sequenced animals, and not shared by other SJL/J mice. These variants were annotated using ANNOVAR [33].

### PBGD assay

Purified erythrocytes and reticulocytes were prepared for assaying PBGD activity as follows. Whole blood was collected by cardiac bleeding, mixed with anticoagulant (1/10 volume 85 mM sodium citrate, 69 mM citrate and 111 mM glucose), immediately centrifuged at 150×*g* for 10 min, and the plasma and buffy coat removed. The retained red cells were then centrifuged through 75% Percoll PBS-buffered solution (800×*g* for 10 min) to remove residual leukocytes. Erythrocytes were depleted of reticulocytes (and the reticulocytes separately retained) by addition of allophycocyanin (APC)-conjugated rat anti-mouse CD71 antibodies (1 μg/mL; clone RI7 217.1.4; eBioscience™) followed by MojoSort™ Mouse anti-APC Nanobeads (1/10 volume; Biolegend^®^) and magnetic affinity separation using a QuadroMACS™ Separator (Miltenyi Biotec). The resulting erythrocyte preparations were confirmed to be free of reticulocytes using flow cytometry (>99.9% pure).

Erythrocytes and reticulocytes were assayed for PBGD by incubating 2 μL cells diluted 50-fold in the assay buffer (50 mM Tris containing 0.2% Triton X-100, 20 mM citric acid, pH 8.2), incubated at 56 °C for 15 min, and finally cooled on ice. Duplicate aliquots (40 μL) of the resulting hemolysates were mixed with 80 μL assay buffer containing 80 μM porphobilinogen (Sigma Aldrich, Castle Hill, Australia) and incubated at 37 °C for 60 min to permit the PBGD-catalyzed generation of hydroxymethylbilane. The reaction was terminated by adding 120 μL 12.5% TCA and incubated for 30 min at ∼23 °C, which results in the non-enzymatic conversion (oxidation) of hydroxymethylbilane to uroporphyrinogen I (URO I). After centrifugation (1000×*g*, 5 min), 150 μL of clarified supernatant was transferred to brown glass vials, stored at 4 °C, and subjected to ultrahigh-performance liquid chromatography (UHPLC) within 72 h to quantify URO I levels. Samples (100 μL) were injected and analyzed in a UHPLC system consisting of a Dionex UltiMate™ 3000 in line with a fluorescence FLD-3400 detector. A Hypersil Gold C18 column (1.9 μm particle size, 50 × 2.1 mm internal diameter; Thermo Fisher Scientific, Australia) was used for elution. The mobile phase consisted of 9% (v/v) acetonitrile in 1 M ammonium acetate–acetic acid buffer, pH 5.16 (solvent A), and 10% acetonitrile (v/v) in methanol (solvent B) [34; 35; 36]. The elution program was 0% B (100% A) for initial 2 min, 0% B (100% A) to 10% B (90% A) for 2–6 min, 10% B (90% A) to 90% B (10% A) for 1 min, and isocratic 90% B for further 2 min. The flow rate was set at 0.8 mL/min and column temperature at 35 °C. Eluents were detected at λ_ex_ = 404 nm and λ_em_ = 618 nm. To quantify standards, a solution containing 400 nmol/L URO I in 12.5% TCA was prepared using a 1 mM stock solution dissolved in 1 M HCl (Sigma Aldrich). The concentration of the standard was calculated using absorbance at λ = 405 nm and ε = 505 × 10^3^ L cm^−1^ mol^−1^ [36]. Standard curves were generated by varying the injection volume and integrating the corresponding peaks. Enzyme activity was calculated as nkat/L cells according to Erlandsen et al 2000 [29].

### *P. falciparum* culture

*P. falciparum* was cultured in O^+^ human erythrocytes at 2.5% hematocrit according to a previously described method [37]. The cell-culture medium (CCM) comprised of RPMI 1640 supplemented with 8.8 mM D-glucose, 22 mM HEPES, 208 nM hypoxanthine (Sigma-Aldrich, Missouri, US), 46.1 nM gentamicin, 2.1 g/L AlbuMAX® I, 2.8 mM L-glutamine (Life Technologies, Australia), and 4.2% (v/v) O^+^ human serum. Cultures were maintained in flasks filled with 1%O_2_/3%CO_2_/96%N_2_ gas mix, and kept in an orbitally shaking incubator at 50 rpm at 37°C. Culture parasitemias were maintained at between 0.5% and 10%, and checked every one to two days by visualizing Giemsa-stained thin blood smear by microscopy. CCM was changed every one to two days by pelleting infected cells at 500x*g* for 5 minutes and suspending them in fresh CCM. Human erythrocytes and sera were provided by the Australian Red Cross Blood Service. Blood was washed thrice using CCM that lacked hypoxanthine, AlbuMAX® I, and human serum, and centrifuged at 2800 RPM for 5 minutes after each wash. The *P. falciparum* strain 3D7 was donated by Robin F. Anders (La Trobe University, Melbourne, Australia). The Pf*Pbgd*-KO and 3D7 wild-type parental lines were gifts from Daniel E. Goldberg (Washington University School of Medicine, St. Louis, MO).

### TUNEL staining and immunofluorescence

Deoxyuridine triphosphate nick-end labeling by the terminal deoxynucleotidyl transferase (TUNEL) labels degraded or sheared DNA, revealing apoptosis or necrosis. TUNEL and cell staining for immunofluorescence microscopy were performed on blood smears from mice infected with *P. chabaudi* infected mice as described [23], but with some modifications. Mouse blood samples were fixed for at least 24 h in Cytofix solution (BD Biosciences) diluted fourfold in PBS. Slides coated with polyethylenimine (0.1% v/v in PBS) were smeared with fixed blood and stained overnight in the reaction mix from the Apo BrdU TUNEL Assay Kit (Molecular Probes, Eugene, OR). TUNEL-labeled DNA was detected using a biotinylated anti-BrdU antibody and streptavidin-Alexa Fluor 594 conjugate (ThermoFisher Scientific, Australia). At least 100 infected cells were counted on each slide. After staining, slides were mounted in SlowFade™ Gold Antifade Mountant with DAPI (ThermoFisher Scientific, Australia) and glass coverslips (#1 from Menzel-Gläser). Slides were examined at room temperature using an inverted Axio Observer fluorescence microscope or a LSM 800 confocal microscope with Airyscan, both using 630× magnification and coupled to a Axiocam 503 monochrome cameras (Zeiss, Australia). All acquired images were acquired and processed by using the Zeiss Zen lite microscope software (Zeiss, Australia).

### Generation of *P. berghei pbgd* knockout

The *P. berghei ANKA pbgd* locus (*PbA_pbgd*, PBANKA_0608000) was targeted as follows. A targeting plasmid, p35/EF5’.K^+^GFP.CAM3’ was modified [40] by replacing the K^+^GFP sequence with firefly luciferase, yielding the vector p35/EF5’.FF.CAM3’. The 500 bp 5’ flanking sequence of *PbA_pbgd* was amplified by PCR using the primers PbA_060800/5’FIF (5’-gacccgcggGATCATTATGGCAAATATTCTGCC) and PbA_060800/5’FIR (5’ gacctgcagTATTGCTATAACTATATTATTCAGTTA), digested with *Sac*II and *Pst*I enzymes and cloned into the corresponding sites of p35/EF5’.FF.CAM3’ to generate p35/5^’^Fl/EF5’.FF.CAM3’. For the 3’ homology arm, the 3’ flanking sequence of *PbA_pbgd* was amplified by PCR using the primers PbA_060800/3’FIF (5’-gacgatatcGGATGAGGCAGCTTTATATTACA) and PbA_060800/3’FIR (5’gacgaattcGTTCAAATCGCGTATTTGAATGTGATGTG), digested with *Eco*RV and *Eco*RI and cloned into the corresponding sites of p35/5^’^Fl/EF5’.FF.CAM3’ to generate p35/5+3^’^Fl/EF5’.FF.CAM3’. The plasmid vector was then linearized with *Sac*II/*Eco*RI and transfected into PbANKA parasites using previously established methods [41]. The transfected parasites were inoculated into a BALB/c naïve mouse by intravenous inoculation and selected for by administering pyrimethamine into the mouse’s drinking water (0.07 mg/mL) for six days. Parasite DNA was extracted and a PCR was conducted to check for successful integration using the following primers PbA_060800_int_F (primer 539, 5’-ATCCGAGCAATATTATCGACAG), hDHFR_R_int (primer 540, 5’-GATGCAGTTTAGCGA-ACCAAC), CAM3’_F_int (primer 541, 5’-GTAAAGGGTTAATTCTTATATGGTC) and Pb060800_int_R (primer 542, 5’-CCATTTTCTCAATACTTACATAG) (Figure S2). The *PbA_pbgd* parasite was single cloned into 6-8 weeks old BALB/c mice by limiting dilution and verified by PCR according to the procedure described previously [42].

### *P. chabaudi* and *P.berghei* infections

Infection experiments used the rodent parasite *P. chabaudi adami* DS (408XZ) and *P. berghei* ANKA. Parasite stocks were prepared by passage through resistant C57BL/6 (*P. chabaudi*) or BALB/c (*P. berghei*) mice as described previously [22]. Experimental mice were infected by IP injection with a dose of 1 × 10^4^ parasitized erythrocytes. Blood-stage parasitemia was determined by microscopically examining thin blood smears obtained by tail bleeding (1 μL per day from days seven to 14 after infection) and staining with 10% Giemsa solution. Percentages of infected erythrocytes were calculated from a minimum of 1,000 erythrocytes counted.

### Study subjects

We studied erythrocytes obtained from four Italian Caucasian AIP patients recruited at the Department of General Medicine, Fondazione IRCC Ca’ Granda, Ospedale Maggiore Policlinico, Milan, Italy. All patients had a confirmed biochemical and genetic diagnosis: two patients were asymptomatic carriers and two experienced typical acute attacks. The biochemical measures associated with the condition are provided in Table S4.

### Collection and preparation of purified erythrocytes

Blood from four patients with AIP and the Italian healthy control subject was collected by venepuncture into 5-mL tubes containing EDTA. These five blood samples were stored at 4°C and shipped together within seven days of collection from the Ospedale Maggiore Policlinico, Milan, Italy to The Australian National University (ANU), Canberra, Australia. Erythrocytes were purified at the ANU by centrifugation (170×*g*, 13 min) to separate them from the plasma and white-cell fractions. The erythrocytes were removed and washed twice in RPMI before being infected with parasites.

### *P. falciparum* growth-inhibition assays

For growth assays, parasites were propagated in normal erythrocytes (from the Australian Red Cross Blood Service), purified using Percoll density-gradient centrifugation [38], and then added to test erythrocytes at a final parasitemia of ∼1%. Parasitemias were determined in samples collected during culturing for up to 72 h. Samples were fixed in Cytofix solution (BD Biosciences) for 24 h, and then washed once in PBS and stained with 5 μg/mL Hoechst 33342 for 5 min. Fluorescence signals were measured using an LSR Fortessa cell analyzer (BD Biosciences). At least 100,000 events were recorded per sample. BD FACS software was used to process and analyze the data.

### Statistical analysis

For malaria survival, statistical significance was determined using the Mantel–Cox test. For other results, two-tailed Students *t*-test or ANOVA were used as indicated. A *p*-value of < 0.05 was considered significant.

## ACKNOWLEDGMENTS

The Authors were supported by an International Macquarie University Research Excellence Scholarship (CBS), the NHMRC (490037, 605524, APP1047090 and APP1066502), the Australian Research Council (DP120100061) and the National Collaborative Research Infrastructure (NCRIS) via the Australian Phenomics Network (APN). Funders had no role in the design of the study and collection, analysis, and interpretation of data or in writing the manuscript. We thank Ceri Flowers for administrative support, Diana Spinelli for recruitment of the AIP patients, Shelley Lampkin for technical assistance, Elinor Hortle for assisting with the cytokine assays, Gavin Symonds and Zeiss Australia for confocal imaging analysis, and the Australian Red Cross Blood Service for red cells and human serum.

## Author Information

Department of Immunology and Infectious Disease, John Curtin School of Medical Research, The Australian National University, Canberra, ACT 2601, Australia.

Cilly Bernardette Schnider, Hao Yang, Lora Starrs, Anna Ehmann, Simon J. Foote, Brendan J. McMorran, Gaetan Burgio

Department of Biomedical Sciences Faculty of Medicine and Health Sciences, Macquarie University, NSW 2109 Australia.

Cilly Bernardette Schnider

Research School of Biology, The Australian National University, Canberra, ACT 2601, Australia. Farid Rahimi

Fondazione IRCCS Ca’ Granda - Ospedale Maggiore Policlinico, Rare Diseases Center, Internal Medicine Unit, Department of Medicine and Medical Specialties, Via Francesco Sforza, 35, 20122 Milan, Italy.

Elena Di Pierro and Giovanna Graziadei

School of Medicine, Deakin University, Warn Ponds, VIC 3216, Australia

Kathryn Matthews and Tania De Koning-Ward

Health and Biosecurity, CSIRO, Sydney, NSW 2113, Australia.

Denis C. Bauer

## Author Contributions

CBS, HY, LS, AE, TKW, KM, FR, and BJM performed experiments and DCB analyzed the exome sequencing data. The project was conceived by BJM, GB and SJF and experiments were designed by CBS, HY, BJM and GB. The blood samples, and the genetic and biochemical analysis of the AIP patients were provided and performed by EDP and GG. The paper was written and the figures were prepared by CBS, EDP, GG, GB and BJM. All authors read and approved the final manuscript.

## Competing interests

The authors declare that they have no competing interests that might be perceived to influence the results and/or discussion reported in this paper.

## Data Availability

All data generated or analysed during this study are included in this article (and its Supplementary Information files)

## SUPPLEMENTAL FIGURE LEGENDS

**Figure S1:**
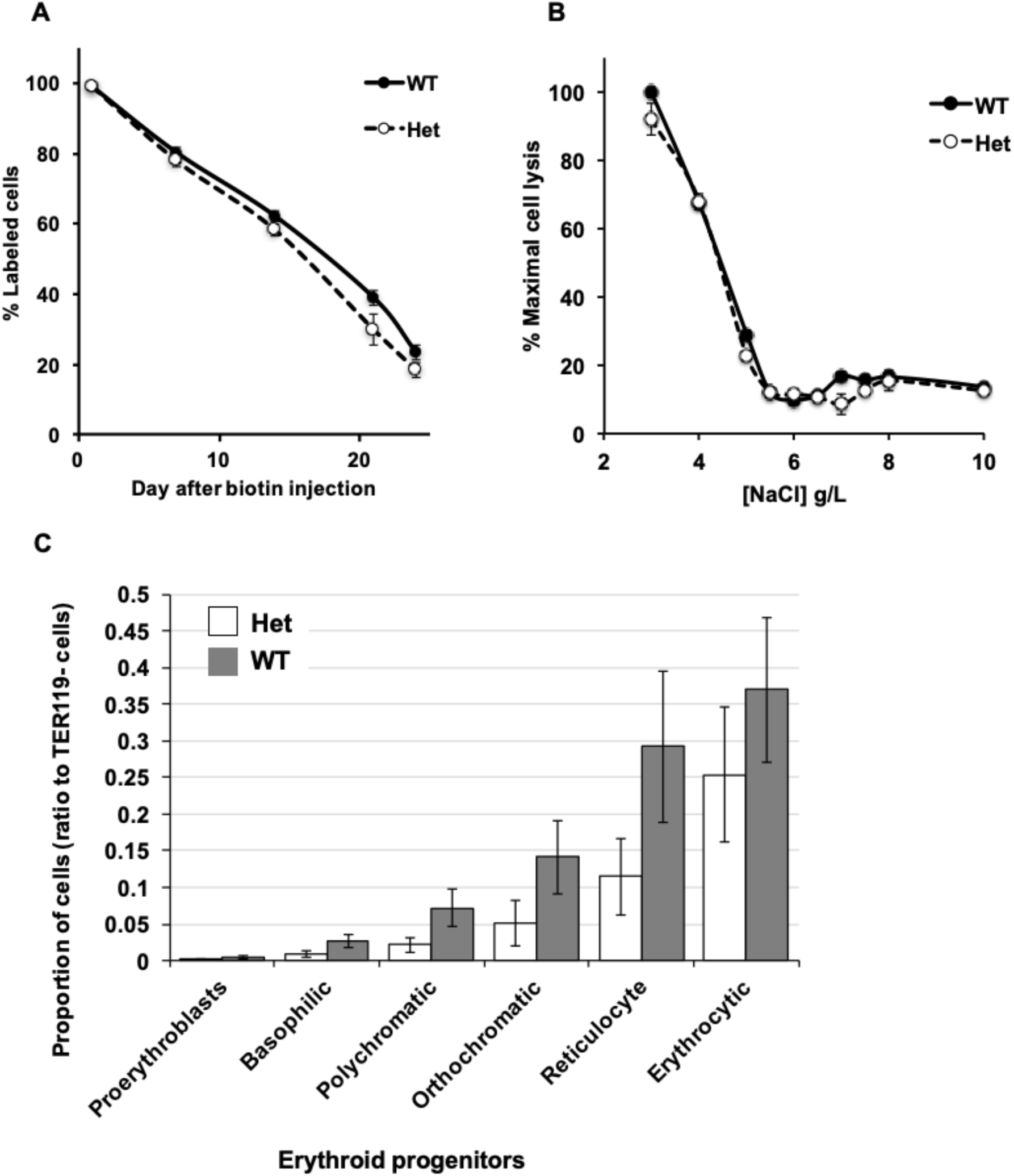
Hematological analysis of the *Pbgd*^*MRI58155*^ mouse line. **(A)** Comparison of erythrocyte life span, (**B)** osmotic strength, and (**C)** bone-marrow erythroid progenitor cells in *Pbgd*^*MRI58155*^ heterozygotes (Het, n = six to nine mice) and wild-type littermates (WT, n = six to seven mice). Error bars represent SEM. No significant differences were observed between groups using the two-tailed Student’s *t*-test assuming equal variance.

**Figure S2:**
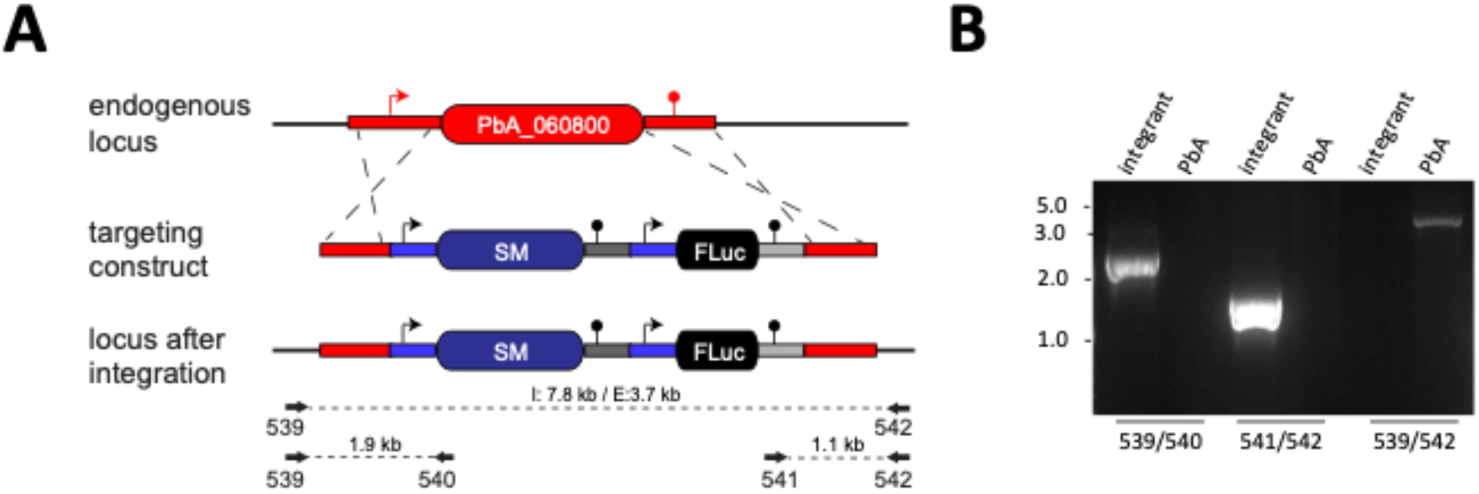
Generation of *Pbgd P. berghei* knockout. **(A)** For targeted gene deletion of the PbANKA_060800, fragments of the 5’ UTR and 3’ UTR, which would serve as targeting regions to drive integration into the endogenous locus, were PCR-amplified from *P. berghei* ANKA genomic DNA and cloned into the p35/EF5’.Fluc.CAM3’ vector. Position of oligonucleotides used for PCR analysis are shown. (**B)** Representation of a gel electrophoresis on the genomic integration of the targeting construct using specific primers spanning the locus. SM, *hDHFR* selectable marker; I, integrant fragment size; E, non-integrant fragment size.

## SUPPLEMENTAL TABLES

**Table S1.** Hematological parameters of wild-type and *Pbgd*^*MRI58155*^ heterozygous mice. Automated full blood analysis on SJL mice (n = 500), wild type (n = 24) and heterozygous (n = 63) mice. RBC = erythrocyte; WBC = white blood cell; MCV = mean corpuscular volume; HCT = haematocrit; % retics = proportion of reticulocytes; HGB = hemoglobin concentration; MCHC = mean corpuscular hemoglobin concentration. Values denote mean ± SEM. ** p < 0.01; *** p < 0.001.

|  | RBC | WBC | MCV | HCT | Retics (%) | HGB (g/L) | MCHC (g/L) |
| --- | --- | --- | --- | --- | --- | --- | --- |
| Het | 10.11 ± 0.17 | 11.09 ± 0.38 | 49.95 ± 0.22*** | 0.50 ± 0.008*** | 5.84 ± 0.26*** | 141.8 ± 2.5*** | 215.5 ± 7.4 |
| WT | 10.25 ± 0.05 | 11.44 ± 0.45 | 52.40 ± 0.23 | 0.54 ± 0.003 | 3.24 ± 0.21 | 153.4 ± 0.8 | 210 ± 6.5 |
| SJL/J | 9.79 ± 0.22 | 11.84 ± 0.50 | 52.61 ± 0.19 | 0.51 ± 0.011 | 3.44 ± 0.033 | 153.84 ± 0.23 | 203.53 ± 7.95 |

**Table S2.** Spleen and liver indices, and non-heme iron content in wild-type and *Pbgd*^*MRI58155*^ heterozygous mice (n = 3 per group). Values denote means ± SD. * p < 0.05.

|  | Spleen index | Liver index | Spleen iron (mg/g) | Liver iron (mg/g) |
| --- | --- | --- | --- | --- |
| Het | 0.477 ± 0.017* | 5.8 ± 0.33 | 15.2 ± 1.2 | 3.59 ± 0.65 |
| WT | 0.408 ± 0.009 | 5.96 ± 0.16 | 11.4 ± 0.2 | 4.58 ± 0.62 |

**Table S3.** Serum cytokine concentrations in wild-type (n = 5) and *Pbgd*^*MRI58155*^ heterozygous (n = 8) mice. All cytokine concentrations are pg/mL plasma. Values denote means ± SD. * p < 0.05.

| | IL-2 | IL-4 | IL-6 | IFNγ | TNFα | IL-10 | IL-17A |
| --- | --- | --- | --- | --- | --- | --- | --- |
| Het | 119 ± 55 | 122 ± 56 | 84 ± 54 | 68 ± 42 | 140 ± 85 | 93 ± 42* | 110 ± 51* |
| WT | 192 ± 51 | 157 ± 56 | 156 ± 87 | 111 ± 53 | 186 ± 92 | 146 ± 34 | 187 ± 53 |

**Table S4.** AIP patient blood biochemistry. * Patients were symptomatic for AIP at the time of blood collection.

| AIP<br>patient<br>ID | ALA/Creatinin<br>e ratio | PBG/Creatinin<br>e ratio | Total urinary<br>porphyrins<br>(µg/L) | % Urine porphyrin types<br>(URO, EPTA, ESA, PENTA,<br>COPRO I, COPRO III) | PBGD<br>activity<br>(pmol/h/mg<br>Hb) |
| --- | --- | --- | --- | --- | --- |
| A | 2.3 | 0.4 | 73 | 4, 1, <1, 10, 37, 48 | 61.1 |
| B* | 18.6 | 11.1 | 179 | 43, 3, 3, 22, 17, 12 | 74.8 |
| C* | 24.4 | 3.4 | 575 | 24, 2, 3, 23, 8, 41 | 56.9 |
| D | 1.5 | 0.3 | 88 | 8, 4, 1, 8, 58, 21 | 57.5 |

## Notes

### Competing Interest Statement

The authors have declared no competing interest.

## REFERENCES

[1] A.C. Allison, Genetic control of resistance to human malaria. Curr Opin Immunol 21 (2009) 499–505.

[2] A.V. Hill, The immunogenetics of resistance to malaria. Proc Assoc Am Physicians 111 (1999) 272–7.

[3] G. Min-Oo, and P. Gros, Erythrocyte variants and the nature of their malaria protective effect. Cell Microbiol 7 (2005) 753–63.

[4] R.L. Nagel, and E.F. Roth, Jr., Malaria and red cell genetic defects. Blood 74 (1989) 1213–21.

[5] Z.Q. Bonday, S. Dhanasekaran, P.N. Rangarajan, and G. Padmanaban, Import of host delta-aminolevulinate dehydratase into the malarial parasite: identification of a new drug target. Nat Med 6 (2000) 898–903.

[6] S. Dhanasekaran, N.R. Chandra, B.K. Chandrasekhar Sagar, P.N. Rangarajan, and G. Padmanaban, Delta-aminolevulinic acid dehydratase from Plasmodium falciparum: indigenous versus imported. J Biol Chem 279 (2004) 6934–42.

[7] S. Koncarevic, P. Rohrbach, M. Deponte, G. Krohne, J.H. Prieto, J. Yates, 3rd, S. Rahlfs, and K. Becker, The malarial parasite Plasmodium falciparum imports the human protein peroxiredoxin 2 for peroxide detoxification. Proc Natl Acad Sci U S A 106 (2009) 13323–8.

[8] A. Sicard, J.P. Semblat, C. Doerig, R. Hamelin, M. Moniatte, D. Dorin-Semblat, J.A. Spicer, A. Srivastava, S. Retzlaff, V. Heussler, A.P. Waters, and C. Doerig, Activation of a PAK-MEK signalling pathway in malaria parasite-infected erythrocytes. Cell Microbiol 13 (2011) 836–45.

[9] S. Varadharajan, B.K. Sagar, P.N. Rangarajan, and G. Padmanaban, Localization of ferrochelatase in Plasmodium falciparum. Biochem J 384 (2004) 429–36.

[10] M. Brizuela, H.M. Huang, C. Smith, G. Burgio, S.J. Foote, and B.J. McMorran, Treatment of erythrocytes with the 2-cys peroxiredoxin inhibitor, Conoidin A, prevents the growth of Plasmodium falciparum and enhances parasite sensitivity to chloroquine. PLoS One 9 (2014) e92411.

[11] C.M. Smith, A. Jerkovic, T.T. Truong, S.J. Foote, J.S. McCarthy, and B.J. McMorran, Griseofulvin impairs intraerythrocytic growth of Plasmodium falciparum through ferrochelatase inhibition but lacks activity in an experimental human infection study. Sci Rep 7 (2017) 41975.

[12] H. Ke, P.A. Sigala, K. Miura, J.M. Morrisey, M.W. Mather, J.R. Crowley, J.P. Henderson, D.E. Goldberg, C.A. Long, and A.B. Vaidya, The heme biosynthesis pathway is essential for Plasmodium falciparum development in mosquito stage but not in blood stages. J Biol Chem 289 (2014) 34827–37.

[13] V.A. Nagaraj, B. Sundaram, N.M. Varadarajan, P.A. Subramani, D.M. Kalappa, S.K. Ghosh, and G. Padmanaban, Malaria parasite-synthesized heme is essential in the mosquito and liver stages and complements host heme in the blood stages of infection. PLoS Pathog 9 (2013) e1003522.

[14] Z. Rizopoulos, K. Matuschewski, and J.M. Haussig, Distinct Prominent Roles for Enzymes of Plasmodium berghei Heme Biosynthesis in Sporozoite and Liver Stage Maturation. Infect Immun 84 (2016) 3252–3262.

[15] P.A. Sigala, J.R. Crowley, J.P. Henderson, and D.E. Goldberg, Deconvoluting heme biosynthesis to target blood-stage malaria parasites. Elife 4 (2015).

[16] Z.Q. Bonday, S. Taketani, P.D. Gupta, and G. Padmanaban, Heme biosynthesis by the malarial parasite. Import of delta-aminolevulinate dehydrase from the host red cell. J Biol Chem 272 (1997) 21839–46.

[17] E.M. Pasini, M. Kirkegaard, P. Mortensen, H.U. Lutz, A.W. Thomas, and M. Mann, In-depth analysis of the membrane and cytosolic proteome of red blood cells. Blood 108 (2006) 791–801.

[18] E.M. Pasini, M. Kirkegaard, P. Mortensen, M. Mann, and A.W. Thomas, Deep-coverage rhesus red blood cell proteome: a first comparison with the human and mouse red blood cell. Blood Transfus 8 Suppl 3 (2010) s126–39.

[19] E.M. Pasini, M. Kirkegaard, D. Salerno, P. Mortensen, M. Mann, and A.W. Thomas, Deep coverage mouse red blood cell proteome: a first comparison with the human red blood cell. Mol Cell Proteomics 7 (2008) 1317–30.

[20] A. Brun, G. Gaudernack, and S. Sandberg, A new method for isolation of reticulocytes: positive selection of human reticulocytes by immunomagnetic separation. Blood 76 (1990) 2397–403.

[21] C.M. Smith, A. Jerkovic, H. Puy, I. Winship, J.C. Deybach, L. Gouya, G. van Dooren, C.D. Goodman, A. Sturm, H. Manceau, G.I. McFadden, P. David, O. Mercereau-Puijalon, G. Burgio, B.J. McMorran, and S.J. Foote, Red cells from ferrochelatase-deficient erythropoietic protoporphyria patients are resistant to growth of malarial parasites. Blood 125 (2015) 534–41.

[22] P.M. Lelliott, S. Lampkin, B.J. McMorran, S.J. Foote, and G. Burgio, A flow cytometric assay to quantify invasion of red blood cells by rodent Plasmodium parasites in vivo. Malar J 13 (2014) 100.

[23] B.J. McMorran, V.M. Marshall, C. de Graaf, K.E. Drysdale, M. Shabbar, G.K. Smyth, J.E. Corbin, W.S. Alexander, and S.J. Foote, Platelets kill intraerythrocytic malarial parasites and mediate survival to infection. Science 323 (2009) 797–800.

[24] H. Puy, L. Gouya, and J.C. Deybach, Porphyrias. Lancet 375 (2010) 924–37.

[25] R.L. Lindberg, C. Porcher, B. Grandchamp, B. Ledermann, K. Burki, S. Brandner, A. Aguzzi, and U.A. Meyer, Porphobilinogen deaminase deficiency in mice causes a neuropathy resembling that of human hepatic porphyria. Nat Genet 12 (1996) 195–9.

[26] S. Clavero, D.F. Bishop, M.E. Haskins, U. Giger, R. Kauppinen, and R.J. Desnick, Feline acute intermittent porphyria: a phenocopy masquerading as an erythropoietic porphyria due to dominant and recessive hydroxymethylbilane synthase mutations. Hum Mol Genet 19 (2010) 584–96.

[27] K. Chen, J. Liu, S. Heck, J.A. Chasis, X. An, and N. Mohandas, Resolving the distinct stages in erythroid differentiation based on dynamic changes in membrane protein expression during erythropoiesis. Proc Natl Acad Sci U S A 106 (2009) 17413–8.

[28] D.C. Bauer, B.J. McMorran, S.J. Foote, and G. Burgio, Genome-wide analysis of chemically induced mutations in mouse in phenotype-driven screens. BMC Genomics 16 (2015) 866.

[29] H. Li, and R. Durbin, Fast and accurate short read alignment with Burrows-Wheeler transform. Bioinformatics 25 (2009) 1754–60.

[30] B. Langmead, and S.L. Salzberg, Fast gapped-read alignment with Bowtie 2. Nat Methods 9 (2012) 357–9.

[31] H. Li, B. Handsaker, A. Wysoker, T. Fennell, J. Ruan, N. Homer, G. Marth, G. Abecasis, R. Durbin, and S. Genome Project Data Processing, The Sequence Alignment/Map format and SAMtools. Bioinformatics 25 (2009) 2078–9.

[32] A. McKenna, M. Hanna, E. Banks, A. Sivachenko, K. Cibulskis, A. Kernytsky, K. Garimella, D. Altshuler, S. Gabriel, M. Daly, and M.A. DePristo, The Genome Analysis Toolkit: a MapReduce framework for analyzing next-generation DNA sequencing data. Genome Res 20 (2010) 1297–303.

[33] K. Wang, M. Li, and H. Hakonarson, ANNOVAR: functional annotation of genetic variants from high-throughput sequencing data. Nucleic Acids Res 38 (2010) e164.

[34] C.M. Benton, C.K. Lim, C. Moniz, and D.J. Jones, Ultra high-performance liquid chromatography of porphyrins in clinical materials: column and mobile phase selection and optimisation. Biomedical chromatography : BMC 26 (2012) 714–9.

[35] C.M. Benton, C.K. Lim, H.J. Ritchie, C. Moniz, and D.J. Jones, Ultra high-performance liquid chromatography of porphyrins. Biomedical chromatography : BMC 26 (2012) 331–7.

[36] E.J. Erlandsen, P.E. Jorgensen, S. Markussen, and A. Brock, Determination of porphobilinogen deaminase activity in human erythrocytes: pertinent factors in obtaining optimal conditions for measurements. Scand J Clin Lab Invest 60 (2000) 627–34.

[37] W. Trager, and J.B. Jensen, Human malaria parasites in continuous culture. Science 193 (1976) 673–5.

[38] E.M. Rivadeneira, M. Wasserman, and C.T. Espinal, Separation and concentration of schizonts of Plasmodium falciparum by Percoll gradients. J Protozool 30 (1983) 367–70.

[39] U.K. Laemmli, Cleavage of structural proteins during the assembly of the head of bacteriophage T4. Nature 227 (1970) 680–5.

[40] S. Haase, E. Hanssen, K. Matthews, M. Kalanon, and T.F. de Koning-Ward, The exported protein PbCP1 localises to cleft-like structures in the rodent malaria parasite Plasmodium berghei. PloS one 8 (2013) e61482.

[41] C.J. Janse, J. Ramesar, and A.P. Waters, High-efficiency transfection and drug selection of genetically transformed blood stages of the rodent malaria parasite Plasmodium berghei. Nature protocols 1 (2006) 346–56.

[42] C.J. Janse, B. Franke-Fayard, G.R. Mair, J. Ramesar, C. Thiel, S. Engelmann, K. Matuschewski, G.J. van Gemert, R.W. Sauerwein, and A.P. Waters, High efficiency transfection of Plasmodium berghei facilitates novel selection procedures. Mol Biochem Parasitol 145 (2006) 60–70.

